# Experimental manipulation of *Heliconius* warning patterns reduces harassment of previously mated females

**DOI:** 10.1101/437525

**Authors:** Richard M. Merrill, Sara Neggazi, Colin R. Morrison, Rachel Crisp, W. Owen McMillan

**Affiliations:** Division of Evolutionary Biology, Ludwig-Maximilians-Universitat, Munich, Germany; Smithsonian Tropical Research Institute, Panama City, Panama

**Keywords:** Sexual conflict, ecological divergence, adaptation, mimicry, magic trait

## Abstract

Why warning patterns are so diverse is an enduring evolutionary problem. Because predators learn to associate particular patterns with unpleasant experiences, an individual’s risk of predation should decrease as the local density of its warning pattern increases. *Heliconius* butterflies, however, are known for their diversity of warning patterns, and the establishment of entirely new phenotypes is difficult to explain under strict frequency-dependent selection. One possibility is that during periods of relaxed selection, drift may allow new variants to rise above a threshold density until mimicry selection takes over. We propose an alternative hypothesis where novel pattern phenotypes arise due to a conflict of interests between the sexes. It is well established that male *Heliconius* use warning patterns as a mating cue. This will likely be beneficial to males as it will increase the efficiency of finding mates. However, already mated females may suffer fitness costs if these cues lead to harassment by males during oviposition or foraging. When constraints imposed by predation are locally relaxed, this could lead to rapid divergence in pattern phenotypes through chase-away sexual selection. To begin to test this hypothesis, we experimentally manipulated the warning patterns of mated *Heliconius erato demaphoon* females and recorded their interactions with conspecific males, and the effect of male presence on laying rate. As predicted, males interacted less with mated females whose red forewing band was blacked-out, as compared to control females whose warning pattern remained intact. We also show that females lay less eggs in the presence of males, but we were unable to detect a significant interaction between warning pattern treatment and the presence of males on female fecundity. Our results suggest that male attraction to conspecific warning patterns, may impose a previously unrecognized cost on *Heliconius* females.

## Introduction

The natural world is characterized by ecological diversity, with organisms employing an apparent endless variety of strategies to survive and reproduce. Natural selection, however, is short-sighted (Darwin 1909; Fisher 1999), presenting an enduring puzzle: If selection can only exploit the best of the immediately available alternative phenotypes, how can novel ecological strategies evolve in organisms that are already well-adapted? This has traditionally been envisaged as the problem of peak shifts across the metaphorical ‘fitness landscape’(Wright 1931). When the environment remains stable, in order to move from one adaptive peak (i.e. local optimum) to another, populations must first transverse a fitness valley, inhabited by intermediate and typically maladaptive phenotypes. To overcome this problem, genetic drift is often invoked as a means by which populations may avoid these fitness valleys (Wright 1931; Coyne and Orr 2004; Mallet 2010).

Sex-specific selection presents an alternative mechanism by which ecological diversity may be promoted (Bonduriansky 2011). Ecological adaptations, including for example, strategies to exploit resources within the environment or avoid predators, are typically – though not always – shared between the sexes. Here, viability selection is normally expected to push these ecological phenotypes towards a shared optimum. Sex-specific selection on-the-other-hand can result in different adaptive optima for the two sexes (Andersson 1994; Arnqvist & Rowe 200). This can result, for example, from requirements imposed on females to produce offspring or the need for males to find receptive mates. The existence of different optima for the two sexes can also lead to sexually antagonistic selection and lead to rapid evolution, even in opposition to viability selection (Arnqvist & Rowe 2005). For example, males may evolve strategies that increase their likelihood of securing mates. If these strategies happen to impose costs on females, females may in turn evolve strategies to circumvent these male tactics, leading to further selection on males and so on. If sexually antagonistic selection involves ecologically relevant traits, it is conceivable it might result in peak shifts across the (viability) fitness landscape.

Since Bates (1862) first described mimicry theory, *Heliconius* butterflies have been the subject of a huge body of work relating to adaptation and speciation (Merrill et al. 2015). These Neotropical butterflies are well known for their bright warning patterns. These are often associated with Müllerian mimicry, where two or more distasteful species converge on the same warning signal to share the cost of educating predators. Because predators learn to associate particular patterns with unpleasant experiences, an individual’s risk of predation decreases as the local density of its warning pattern increases. Warning patterns are predicted to be an important ecological adaptation in *Heliconius.* Three lines of evidence support this claim: First, evidence for learning based on previous experience of birds studied both in captivity (Chai and Srygley 1990; Merrill et al. 2012) and in the wild (Langham 2004); second, higher recapture rates of released butterflies that match locally more abundant warning patterns (Benson 1972; Mallet and Barton 1989; Kapan 2001); and finally, lower attack rates on artificial butterflies matching local co-mimics (Finkbeiner et al. 2012; Merrill et al. 2012; Chouteau et al. 2016). Selection coefficients for local warning patterns are strong (Mallet et al. 1990). As such, local patterns represent a considerable fitness peak in the adaptive landscape, which has presumably led to the observed and widespread convergence of prey species sharing a habitat.

Despite strong positive frequency dependent selection, *Heliconius* butterflies exhibit a massive diversity of alternative warning patterns (Bates 1862; Merrill et al. 2015). Individual *Heliconius* species often vary in warning pattern across their range, leading to distinct geographical colour pattern races (the most prominent being the parallel radiations of *H. erato* and *H. melpomene*); in some cases, such as in *H. numata*, *H. doris*, and occasionally *H. cydno*, warning pattern polymorphisms also exist within single geographical populations. In addition, multiple warning patterns (involving both individual species, or two or more taxa forming mimicry rings) frequently coexist within a single geographical community. A given *Heliconius* species may then join many distinct mimicry rings according to local context, leading to the well-documented mosaic of warning patterns observed across the Neotropics (Brown 1976). Spatial variation in local predator and prey communities, as well as perhaps the local signaling environment, shapes a rugged adaptive landscape with separate fitness peaks, and is therefore crucial to the diversification of warning signals.

Nevertheless, the establishment of entirely new phenotypes remains problematic under strict frequency dependent selection. One possibility is that during periods of relaxed selection, drift may allow new variants to rise above a threshold density until mimicry selection takes over (Mallet and Joron 1999; Sherratt 2006; Mallet 2010). Another possibility is that a model in which predators only learn to avoid unpalatable prey only after sampling a fixed number is overly simplistic. Specifically, if predators are neophobic and generally avoid prey with unfamiliar phenotypes, novel signaling phenotypes might be favoured when they are rare (Aubier and Sherratt 2015). Here we propose an additional factor that may contribute to the origin of novel warning patterns – specifically, that females may benefit from uncommon wing patterns because they will lead to reduced harassment by males, which use colour pattern as a mating cue.

Jocelyn Crane (1955) first demonstrated that the bright warning patterns of *Heliconius* stimulate male courtship in the 1950s. Since then, numerous insectary experiments using both live females and artificial butterflies have repeatedly shown that male *Heliconius* almost always prefer ‘females’ that share their own wing pattern phenotype over that of other conspecific races or closely related species (Jiggins et al. 2001, 2004; Kronforst et al. 2006; Melo et al. 2009; Merrill et al. 2011, 2014). It seems likely that competition between males drives local preferences as the ability to efficiently locate potential mates within a visually complex environment would be beneficial. However, already mated females may suffer fitness costs if these cues lead to harassment by males during oviposition or foraging. These costs would be augmented by the fact that although individual *Heliconius* are long lived (up to 6 months), female re-mating is a rare event (Walters et al. 2012). To begin to test this hypothesis, we experimentally manipulated the warning pattens of mated *Heliconius erato* females and recorded i) their interactions with conspecific males, and ii) the effect of male presence on laying rate.

## Materials and Methods

### BUTTERFLY COLLECTION AND MAINTENANCE

*Heliconius erato demaphoon* were collected in Gamboa and the nearby Soberania National Park, Panama. Females were maintained in a 2 × 2 × 2m cage with ~10% sugar solution, pollen source and *Passiflora biflora* host plants before the experiments, with other females but no males. Males, collected from the same area, but numbered on their forewing to enable individual identification, were maintained in a similar cage. All experiments were performed between November 2014 and October 2016 in the Smithsonian Tropical Research Institute insectaries in Gamboa, Panama.

### MALE HARRASSMENT EXPERIMENTS

Females were introduced individually into an experimental cage with ~10% sugar solution, a pollen source and a single *P. biflora* host-plant, and then left for 48 hours. Individual *P. biflora* host-plants were not re-used in subsequent trials with different females. The initial 48-hour period of cage-acclimatization was followed by two 48-hour experimental periods followed. Three *H. erato* males were introduced into the experimental cage with the focal female during either the first or second of these experimental periods. Thus, each female experienced 48 hours with, and 48 hours without conspecific males in a randomized order. Although individual males were re-used in subsequent trials, individual identification numbers allowed us to ensure no two females experienced the same combination of males.

The first 15 females included in our experiment (between 19^th^ November 2014 and 14^th^ January 2015) underwent no colour pattern treatment (‘preliminary experiment’). At the start of each trial, all subsequent females were assigned to an experimental warning pattern treatment (‘main experiment’): i) Disruption of the warning pattern by painting over the entire red area of the dorsal side of the forewings with a black Copic™ marker; or one of two control treatments ii) painting over the entire red area of the dorsal side of the forewings with a colourless marker, or iii) handling but no marker. Only females from the main experiment were included in analyses testing the effects of warning pattern treatment; to maximize experimental power, females from both the preliminary and main experiments were included in analyses that do not consider warning pattern treatment.

We carried out observations for a subset of females, recording the time and duration of male-female interactions (hovering ‘courtship’ and chasing). Due to logistic necessities the total observation time differed between females, largely relating to whether observations were carried out on just one or both days in which females were housed with males. For subsequent analysis we consequently used i) the time until first interaction, and ii) proportion of total observation period in which interactions occurred. For females that were observed on two days, and where no interactions occurred on the first day, time until first interaction was calculated as the total length of first observation period plus the time until interaction on the second day. We also recorded the number of eggs laid by females every 24 hours. Because laying eggs is often a sign of female condition, we excluded two females from subsequent analysis that did not lay eggs during either of the 48-hour experimental periods (though their inclusion does not quantitatively affect our results).

### FEMALE CHOICE EXPERIMENT

We tested for an effect of intact vs disrupted warning pattern on female preference in choice trials. Individuals for these experiments were raised from eggs. Caterpillars were fed individually in pots on *Passiflora* leaves until 4^th^ instar, at which point they were transferred into small (~30cm × ~45cm) cages and fed in groups on potted host plants until pupation. Males and females were separated into different cages and marked on the forewing patch with the date of eclosion. One day old virgin females were introduced into a cage with two males that were at least 10 days old, and no older than 30 days, between 0800 and 1000 hrs. The previous day, males had been assigned the one of two treatments: i) the experimental male had the red forewing and yellow hindwing scales disrupted by a black Sharpie™; ii) the control male had black Sharpie™ marker applied over black scales so that his red bar remained intact. Males used in experiments had never experienced a female. The experimental cage was checked for mating every hour, during daylight hours until 1500hrs the follow day, and mating outcome (experimental, control or none) was recorded.

### STATISTICAL ANALYSES

To measure the cumulative probabilities of avoiding male ‘harassment’ for experimental (warning pattern disrupted) and control females we used ‘survival analysis’. We adopted this approach because it allowed us to consider the rate at which male-female interactions occur rather than simply whether or not they occur, and also because it allows for different lengths of total observation by censoring data. This approach allowed us to estimate the comparative risk of harassment by all three experimental treatments over time.

To test for the effects of males on the number of eggs laid by focal females, we used generalized linear mixed models (GLMMs, implemented using the R package lme4), with Poisson errors. Male presence, warning pattern treatment and their interaction were included as fixed effects in our fully saturated model. This was simplified in a stepwise manner testing for reduction in deviance with likelihood ratio tests. Individual female id was included as a random factor in all models to account for the repeated measures design of our experiment. Our prediction was that any differences in eggs laid as a result of male attraction to warning patterns would be observed as a significant interaction term between male presence and warning pattern treatment. All statistical analyses were carried out in R (https://www.r-project.org).

## Results

### WARNING PATTERNS INFLUENCE MALE HARASSMENT OF MATED FEMALES

Of individuals that laid eggs during the main experiment, we carried out observations for 58 individual females, including 30 (‘experimental’) butterflies with disrupted warning patterns, and 28 (‘control’) butterflies subjected to one of the two control treatments (14 of each). Females with intact warning patterns were harassed more quickly than those with disrupted warning patterns (Figure 1a). More specifically, the probability of avoiding male harassment declined more quickly for control females, with intact warning patterns, compared to experimental females with disrupted wing pattern (2ΔlnL = 7.18, d.f. = 1, *P* = 0.007). There was no significant reduction in deviance gained by separating the two control treatments (2ΔlnL ≪ 0.01, d.f. = 1, *P* = 0.99). Overall, male-female interactions were observed more often in trials involving control females without disrupted warning patterns (χ^2^ = 4.35, d.f. = 1, *P* = 0.037; not correcting for length of observation), despite slightly shorter observations for control butterflies (mean = 48mins, median = 60mins, range = 16-62mins) than for experimental butterflies (mean = 51mins, median = 60mins, range = 30-120mins).

**Figure 1.**
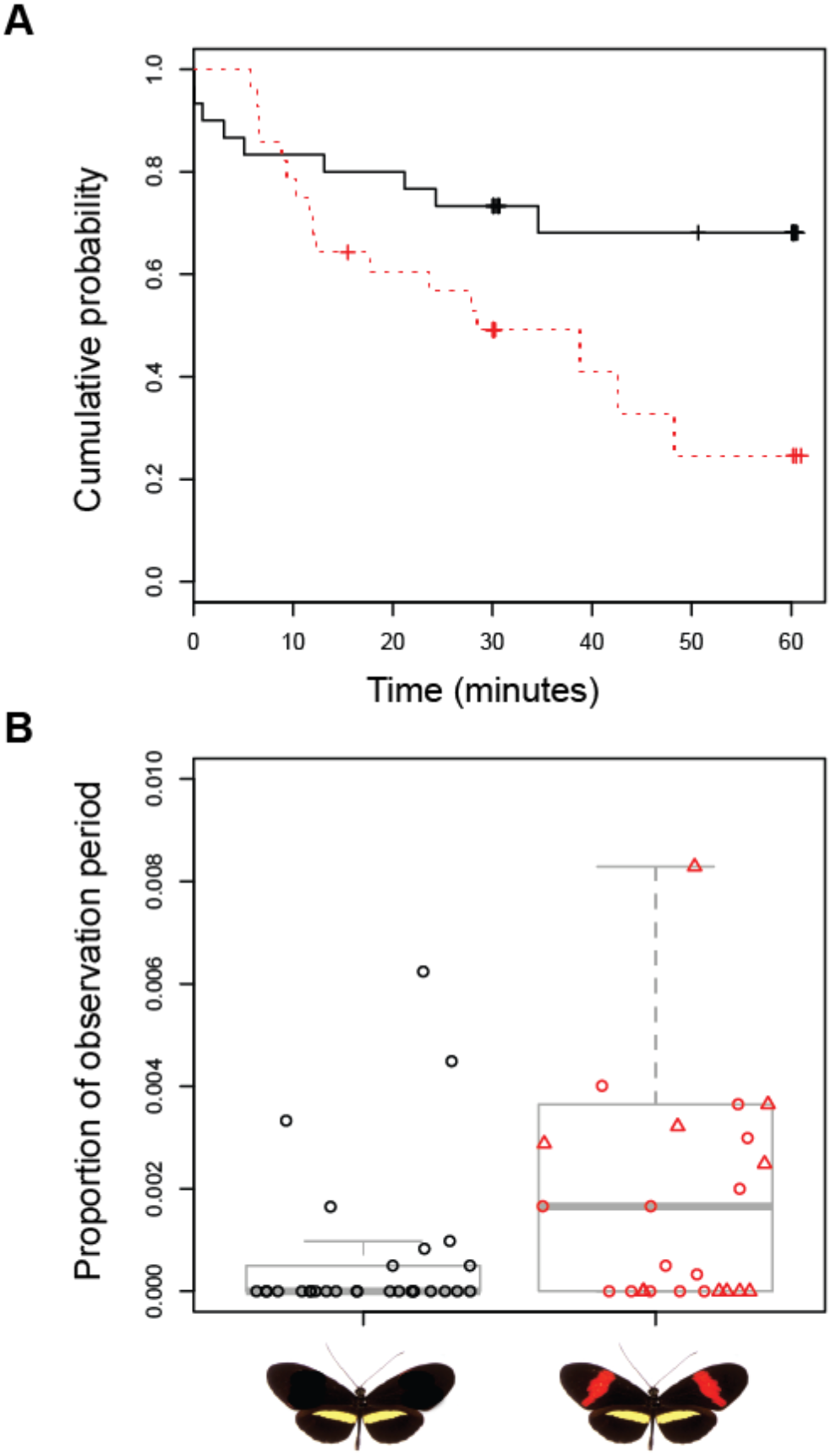
Males harass mated females with intact warning patterns more quickly (A) and for more time (B) than females with disrupted warning patterns. A. Kaplain-Mair plot showing decline in cumulative probability of avoiding male harassment for females with disrupted warning patterns (solid, black) and females with intact warning patterns (dashed, red). B. Boxplots of the proportion of total observation time males spend harassing females with disrupted warning patterns (left, black) and females with intact warning patterns (right, red). Red circles and triangles represent the two control treatments, painting over the entire red area of the dorsal side of the forewings with a colourless marker, or handling but no marker, respectively.

**Figure 2.**
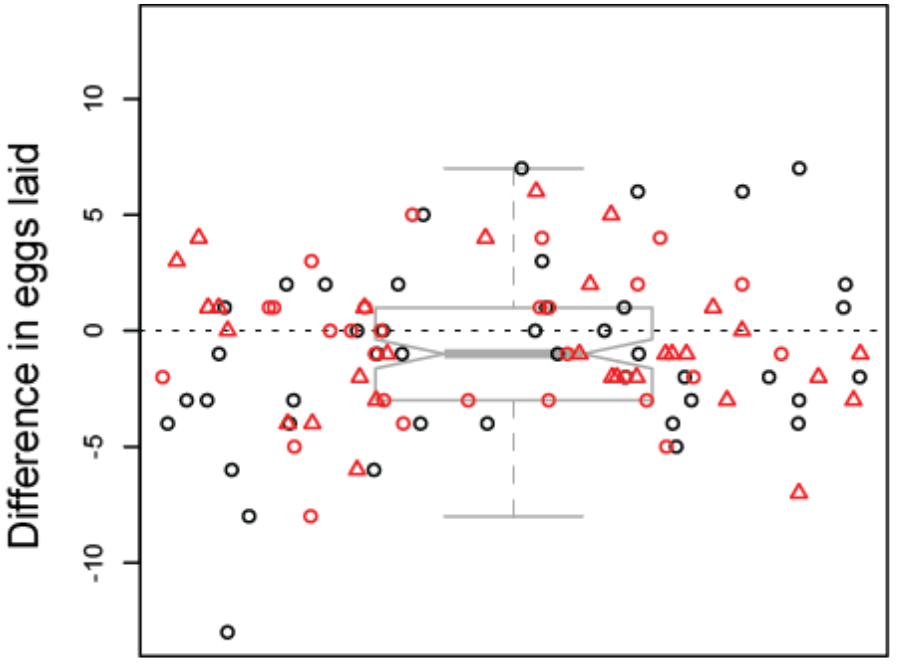
Females lay fewer eggs in the presence of males. Black circles represent females whose warning patterns were experimentally disrupted. Red shapes represent control females with intact warning patterns (circles = painting over the entire red area of the dorsal side of the forewings with a colourless marker; triangles = handling but no marker).

Males also spent less time interacting with experimental females (Wilcoxon rank sum test: *n* = 58, W = 273.5, *P* ~ 0.013; figure 1b). Again, there was no significant difference between the two control treatments (Wilcoxon rank sum test: *n* = 30, *W* = 107, *P* ~ 0.845). Excluding a female that was harassed by males for more than 30% of the observation period did not quantitatively affect the results (Wilcoxon rank sum test: *n* = 59, *W* = 282.5, *P* ~ 0.01).

### MALE PRESENCE REDUCES THE NUMBER OF EGGS LAID BY FEMALES

There was considerable variation in the number of eggs laid both between experimental periods (i.e males present and males absent) for individual females, as well as between females (figure 2). Nonetheless, overall, females included in our experiment laid fewer eggs in the presence of males (paired t-test: *n* = 101, *t* = −2.305, d.f. = 100, *P* = 0.023; figure 2). This effect was subtle and, on average, over a two-day period, females laid ~0.8 fewer eggs in the presence of males. However, overall this represents ~10% decrease in eggs laid compared to when males where absent. We found no evidence of an interaction between the presence of males and warning pattern treatment on the number of eggs laid (2ΔlnL = 2.067, d.f. = 2, *P* = 0.356).

### MANIPULATION OF MALE WARNING PATTERN DOES NOT INFLUENCE FEMALE CHOICE

We found no difference in mating success between males whose warning pattern had been disrupted, compared to controls (Exact binomial test, *n* = 25, *P* ~ 1). Across 47 trials, 22 did not result in mating. In the remaining trails, females mated with the control male in 12 trials, and with the experimental (disrupted warning pattern) in 13 trials: The probability of mating with males with intact warning patterns was 0.48 (0.28 − 0.69), strongly suggesting that our warning pattern treatment had little influence on female choice.

## Discussion

The warning patterns of *Heliconius* butterflies have become a textbook example of natural selection (e.g.(Barton et al. 2007)), but the origins of their considerable diversity remain problematic. Here, we provide some experimental evidence that sex-specific selection provides a possible mechanism by which novel warning patterns may initially increase in frequency. Indeed, mated females with intact warning patterns were harassed by conspecific males more quickly and for longer periods, compared to controls with experimentally disrupted patterns. *Heliconius erato* females mate very early after eclosion, and despite a long adult life, almost invariably, do not re-mate (Walters et al. 2012). As such, this disturbance by males would have no obvious benefits to previously mated females. In fact, our experiments highlight a potential cost and females in our experimental trials laid fewer eggs in the presence of males. Within the context of our experiment, this effect was relatively subtle, with an average reduction on of just ~0.8 eggs over the course of two days. However, within the context of female adult life-span of several months, and given that females only lay a handful of eggs each day, and large population sizes, this could potentially have large effects on the evolution of emerging warning patterns.

Although we have shown that the presence of males can reduce the short-term fecundity of females, we were unable to experimentally demonstrate a link between colour pattern and a reduction in the number of eggs laid. In hindsight, this is perhaps not surprising. Even in the absence of males, there was considerable variation between days in the number of eggs laid by individual females. Given the complex behavioral ‘decisions’ involved in egg laying, this presumably results from environmental factors that we are unlikely to be able to identify, and therefore control. We included a relatively large number of females (n = 87) in our analysis of warning pattern manipulation and male presence (particularly for behavioural experiments with *Heliconius* butterflies); nevertheless, unless effect sizes were considerable, power to detect a significant interaction between colour pattern treatment and male presence would remain low without much larger, and likely unachievable, sample sizes.

Although they stimulate male courtship, there is currently little evidence that warning patterns themselves are under sexual selection. Mating is costly for male *Heliconius*, as a result of the transfer of a large nutritious spermataphore and this may lead to the evolution of choosy males: However, given their long adult life-span and infrequent female re-mating, it is unlikely that receptive males (or rather their sperm) are ever a limited resource (Sexual Selection n.d.). Indeed, from a sample of 251 wild caught *H. erato* females, Walters et al. (2012) recorded just a single virgin; broadly similar results are reported for females from 27 other *Heliconius* species, albeit with smaller sample sizes. Competition among females to secure mates would be an unlikely scenario in *Heliconius*, and as such, male preferences are unlikely to impose strong sexual selection on female traits (such as wing pattern).

Recent work has experimentally demonstrated female preferences in *Heliconius* (Southcott and Kronforst, 2017.; Darragh et al. 2017), but there is currently little evidence that females use colour pattern to distinguish between males, though it is not inconceivable (see also (Chouteau et al. 2017)). The results of our female choice trails support this. However, it should be noted that experimental disruption of warning pattern is not the same as testing for rejection of the ‘wrong’ pattern. Here we were testing for the importance of an intraspecific signal. It remains possible that females may use colours of interspecific males (or males belonging to a different colour pattern morphs) as a cue for rejection. Indeed, because interspecific, non-mimietic hybrids are more likely to be predated (Merrill et al. 2012), speciation theory strongly predicts that reinforcement selection will drive the evolution of female rejection based on heterospecific colour patterns as a cue (Felsenstein 1981; Butlin and Smadja 2018). Traits under sexual selection imposed by female choice, and those used to discriminate against unsuitable partners due to reinforcement selection will not necessarily be the same.

Rather than the target of sexual selection, *Heliconius* warning patterns likely impose sexual selection on male preferences by favouring males more strongly attracted to local morphs. Within a complex visual environment, these males will be those most likely to find receptive unmated females, thereby increasing their reproductive success. In the terminology of Arqvist and Rowe (p.17, 2004, and others before them) warning patterns might be thought of as female ‘preference’ traits in that they “bias … fertilization success toward certain male phenotypes” (in this case particular behavioral phenotypes). Considering the sparsity of receptive female in the environment, benefits gained by securing access to females likely outweigh costs associated with attraction to previously mated conspecific females, or even those of co-mimics (Estrada and Jiggins 2008).

A role for sexual sex specific selection driving divergence in primarily ecological traits has been suggested previously (Bonduriansky 2011). In poison the frog, *Oophaga pumelo*, for example, it seems that sexual selection, due to female preferences for bright colours, has contributed to inter-population differences in the frogs’ warning signal (Maan and Cummings 2009). Similarly, evidence suggests that male harassment drives phenotypic diversity in wing colour in damselflies (Svensson et al. 2005). Although there seems to be little evidence that these colour morphs are ‘ecologically relevant’ (*i.e.* affecting an individual’s survival due to interaction with heterospecific individuals, or the abiotic environment), diversity appears to enhance population performance more generally by reducing overall fitness costs to females from sexual conflict (Takahashi et al. 2014). Among *Papilio* butterflies, which are often female-limited Batesian mimics, it has been suggested that non-mimetic ‘male-like’ forms exist to avoid unwanted attention of males; and in *Papilio dardanus* males do indeed prefer to approach mimetic over non-mimetic (male-like) females (Cook et al. 1994). Our study contributes to this body of work by explicitly testing for fitness effects resulting from sexual conflict relating to an ecological trait.

In conclusion, divergence in female (and male) wing patterns in *Heliconius* is driven primarily by strong selection for mimicry, and is likely to impose divergent sexual selection on male preferences to improve their ability to find receptive females. Female re-mating is a rare event (Walters et al. 2012), and males must compete to find virgin females within a visually complex environment (Merrill et al. 2015). As such, divergence in *Heliconius* male preferences is likely driven by initial divergence in female wing pattern. Given our results, and what we know of *Heliconius* biology, it is plausible, but far from proven, that sexual conflict may have played an additional role in the diversification of warning patterns in these butterflies.

## AUTHOR CONTRIBUTIONS

RMM conceived and designed the main experiment, analysed the data and wrote the paper. RMM, SN and RC collected the data. CRM designed and collected the data for the female choice experiment. WOM provided intellectual and logistical support and helped draft the paper. All authors commented and approved the manuscript.

## ACKNOWLEDGEMENTS

We thank the Smithsonian Tropical Research Institute for support and the Ministerio del Ambiente for permission to collect butterflies in Panama. We are grateful to Chris Jiggins for valuable discussion, and for not harassing RMM to complete other projects first. RMM is funded by a DFG Emmy Noether Fellowship and SN was supported by a British Ecological Society Research Grant awarded to RMM.

## DATA ACHIVING

Scripts for analysis and raw data used in analyses are available online: DRYAD REPOSITORY XXX.

